# Enabling strong acetogenic growth on CO_2_ and H_2_: H_2_ solubility limits *Clostridium ljungdahlii* growth on CO_2_ and H_2_

**DOI:** 10.1101/2025.10.22.683911

**Authors:** Noah B. Willis, Paige A. Bastek, Eleftherios T. Papoutsakis

## Abstract

Due to their ability to convert CO_2_, a greenhouse gas, into useful products, certain acetogenic bacterial species, such as *Clostridium ljungdahlii*, have been proposed as promising platform strains for renewable, carbon-negative chemical production. *C. ljungdahlii*, and similar acetogens, grows slowly and produce primarily acetate when grown on CO_2_ with H_2_ as the electron donor, but it grows quickly and can produce ethanol when grown on higher energy substrates, notably CO or fructose. Here, by utilizing different mixing strategies (and notably the first time use of roller bottles) to modulate the volumetric gas interfacial mass transfer coefficient (k_L_a), we show that, under both mixotrophic (sugar and gas utilization) and autotrophic conditions, *C. ljungdahlii* growth and CO_2_ fixation are primarily electron-limited due to the low solubility of H_2_ relative to CO and CO_2_. We demonstrate that, with sufficiently high H_2_ mass transfer, *C. ljungdahlii* can grow at similar high rates using CO_2_ as its sole carbon source compared to CO or fructose, a finding with significant implications for the use of acetogens in CO_2_-negative biomanufacturing, especially because at least 50% of CO used is oxidized and released as CO_2_. We also show that accumulation of fructose inhibits CO_2_ utilization by *C. ljungdahlii* under mixotrophic growth conditions, suggesting that a non-classical “catabolite repression” by fructose inhibits CO_2_ utilization.

## 1. INTRODUCTION

The term “acetogen” refers to any member of the group of anaerobic bacteria capable of growing on and producing acetate (as well as other chemicals) from a diverse range of C1 substrates (most notably CO_2_) via the Wood-Ljungdahl pathway (WLP) (1). Acetogen-catalyzed conversion of CO_2_, H_2_, and/or CO to soluble products (“gas fermentation”) has been proposed as a scalable approach for renewable, carbon-negative chemical production (2). Though acetogens are a highly diverse group of bacteria, which can be found in at least 23 different bacterial genera, the three best studied acetogenic bacteria are *Moorella thermoacetica* (a thermophile), *Acetobacterium woodii*, and *Clostridium ljungdahlii* (3). Likely because it is both mesophilic and can grow on CO (*A. woodii* cannot use CO as a sole carbon source (4)), *C. ljungdahlii* (and its close relative *Clostridium autoethanogenum*) has received the bulk of research attention as a potential workhorse organism for gas fermentation.

Using *C. ljungdahlii* or *C. autoethanogenum* with CO or syngas (mixtures of CO, H_2_, and CO_2_) as the gaseous substrates, several publications have demonstrated production of important commodity chemicals, such as ethanol (5, 6), isopropanol (7, 8), acetone (8, 9), and 2,3-butanediol(10). Robust mixotrophic performance, where fructose is co-utilized with syngas, has also been demonstrated with *C. ljungdahlii* (9). Industrially, the company Lanzatech has successfully commercialized a process using *C. autoethanogenum* as a biocatalyst to convert syngas, generated as a byproduct from the steel industry, into ethanol (11).

However, *C. ljungdahlii* and *C. autoethanogenum* do not grow nearly as well when they use CO_2_, as opposed to CO, as the sole carbon source. In one striking example, in a study which used a bioreactor to control pH and total gas pressure, *C. autoethanogenum* accumulated up to an OD_600_ of 8.4 under an 80/20 mixture of CO/CO_2_, compared to an OD_600_ of only 1.6 under a 60/40 mixture of H_2_/CO_2_ (12). Another study, which compared *C. autoethanogenum* cell extract enzyme activities when grown on CO/CO_2_ vs. CO_2_/H_2_, specifically described growing *C. autoethanogenum* on CO_2_ and H_2_ as “extremely difficult” and used a retentostat (a continuous perfusion culture with complete cell retention, whereby the medium is replenished continuously) to prepare the cells grown on CO_2_ and H_2_ (doubling time of 33 hr), as opposed to a standard chemostat used to prepare the cells grown on CO (doubling time of 9 hr) (13).

Slow acetogenic growth on CO_2_ represents a significant drawback of gas fermentation since one of the major selling points is the proposed ability to mitigate climate change via assimilation of CO_2_ to valuable products. Not only do clostridial acetogens prefer CO to CO_2_, but, when grown on CO, both *C. ljungdahlii* and *C. autoethanogenum* oxidize large fractions of CO to CO_2_. When ethanol, 2,3-butanediol, and acetate are produced using pure CO, 67%, 64%, and 50% of carbon from CO is released as CO_2_, respectively (12). Theoretically, some of this CO_2_ can be re-assimilated by adding H_2_ to the process, but, when grown on CO_2_ and H_2_, *C. ljungdahlii* and *C. autoethanogenum* produce acetate as the major product (12, 13), as opposed to the wider range of more reduced products, such as alcohols, which can be synthesized from CO. These limitations of acetogenic gas fermentation using CO_2_ relative to CO have previously been hypothesized to be due to the lower energy yield of CO_2_ from the WLP relative to CO (12, 14) and/or carbon catabolite repression of CO_2_ utilization by other carbon sources (15).

Here, we explore several strategies to improve the growth and CO_2_ utilization of *C. ljungdahlii* under mixotrophic (fructose, CO_2_, and H_2_) and autotrophic (only CO_2_ and H_2_) growth conditions. Our results demonstrated that, when grown mixotrophically, fructose inhibits CO_2_ utilization by*C. ljungdahlii*, likely due to some form of catabolite repression (though not necessarily classic “carbon catabolite repression”, which typically refers to hierarchical utilization of different sugar substrates). When fructose accumulation was minimized using fed-batch cultures, CO_2_ utilization increased, but these increases reached a point of diminishing returns at higher *C. ljungdahlii* cell densities. Using autotrophic growth experiments, we show that growth of *C. ljungdahlii* on CO_2_ is H_2_ transport-limited and conclude that the poor growth of *C. ljungdahlii* (and similar acetogens) on CO_2_ (relative to CO) is due almost entirely to the 9.5-fold lower Henry’s Law constant of H_2_ compared to CO, not bioenergetics or catabolite repression by other carbon sources. We identify a potential reason why this nearly full order-of-magnitude difference in solubility between H_2_ and CO has been overlooked in the literature and discuss several promising methods to overcome H_2_-transfer limitation and enable scalable, CO_2_-negative fermentations.

## 2. MATERIALS AND METHODS

### Plasmid and strain construction

Construction of the *C. ljungdahlii* p100ptaHALO strain used throughout this study has been previously described (16).

### Microorganisms and growth media

*C. ljungdahlii* p100ptaHALO cultures were grown in Turbo CGM media (17) without glucose and with 30 mM supplemental butyrate (“Turbo CGMB”) (18, 19) and 100 μg/ml clarithromycin for all fermentation experiments. To initiate *C. ljungdahlii* fermentations, a frozen stock was inoculated into 20 mL of YTF medium (20) supplemented with3.5 g/L arginine and 25 g/L MES (“YTAF-MES”) (21) with 100 μg/ml clarithromycin in a 160 mL serum bottle which had been pre-flushed for 2 min with an 80% H_2_, 20% CO_2_ gas mix. The serum bottle was pressurized to 20 psig [137.9 kPa (gauge); 239.3 kPa (abs)] with the 80/20 (H_2_/CO_2_) gas mixture and grown for 24 hours at 37°C on a rotating incubator platform (Southwest Science) at 90 rpm. This pre-culture was used to inoculate pre-flushed serum bottles with Turbo CGMB media with 5 g/L fructose and 100 μg/ml clarithromycin. These pre-cultures were grown to exponential phase (OD_600_ of 0.6-1.2) and used to inoculate the various fermentation experiments. For *C. ljungdahlii*, 1 OD_600_ = 0.377 gDCW·L^-1^ (22). Throughout this study, we report *C. ljungdahlii* biomass in units of OD_600_ since these values represent a direct measurement and will likely be best understood by the reader.

### Fed-batch fructose feeding to serum bottle cocultures

Fed-batch feeding of fructose to serum-bottle cocultures was accomplished using a 1600 Series Six Channel Syring Pump (New Era Pump Systems Inc., USA). For all fed-batch experiments, concentrated fructose feed solution (Turbo CGMB with varied fructose concentrations) was delivered at 0.11 mL per hour to 20 mL of culture volume in 160 mL serum bottles. The fructose feed rate was adjusted by changing the concentration of the fructose stock solution (Table 1). To supply sufficient CO_2_ and H_2_ without over-pressuring the culture (which can inhibit or damage the syringe pump), the headspace of each culture-containing 160 mL serum bottle was connected, via vinyl tubing with luer lock connections, to the headspace of a 500 mL serum bottle. Each paired 500 mL serum bottle was flushed and pressurized to 10 psig (the maximum pressure tolerable by the syringe pump) with 80/20 H_2_/CO_2_ gas mix at the beginning of fermentation. The syringe pump was placed on the upper shelf of a New Brunswick Scientific Innova 4200 Incubator Shaker. The serum bottles were placed below the syringe pump on the shaker platform, incubated at 37 °C, and agitated at 90 RPM.

**Table 1.** Fructose feed rates used for C.ljungdahlii fed-batch fructose feeding experiments.

| Feed Condition | Target Initial Cell Density (OD<sub>600</sub>) | Volumetric Feed Rate (mL·hr<sup>-1</sup>) | Feed Fructose Concentration (mM) | Fructose Feed Rate (mM·hr<sup>-1</sup>) | Initial Cell-Specific Fructose Feed Rate (mM·hr<sup>-1</sup>·OD<sub>600</sub><sup>-1</sup>) |
| --- | --- | --- | --- | --- | --- |
| A | 1 | 0.11 | 135 | 0.75 | 0.75 |
| B | 1 | 0.11 | 68 | 0.25 | 0.25 |
| C | 1 | 0.11 | 34 | 0.13 | 0.13 |
| D | 1 | 0.11 | 17 | 0.083 | 0.083 |
| E | 2 | 0.11 | 34 | 0.13 | 0.083 |
| F | 4 | 0.11 | 68 | 0.25 | 0.083 |

### Roller serum bottle cocultures

Roller bottle *C. ljungdahlii* cultures were performed by preparing 20 mL of culture volume in a 160 mL or 500 mL serum bottle (as described in the Results) and placing the bottle on a Scilogex SCI-T6-S Analog tube roller at the highest speed setting. The tube roller was placed on the upper shelf of a New Brunswick Scientific Innova 4200 Incubator Shaker maintained at 37 °C.

### Metabolite analysis

High pressure liquid chromatography (HPLC) was used to quantify supernatant sugar and solvent concentrations as previously reported (9, 23, 24).

### Identifying the most applicable Henry’s Law constant of carbon monoxide (k_H,CO_) for gas fermentation

On the National Institute of Standards and Technology (NIST) WebBook for CO, six values are reported for the Henry’s Law constant of CO. Five of these values are in the range of 0.82-0.99 mM·bar^-1^, the same order of magnitude as the Henry’s Law constant for H_2_ (0.78 mM·bar^-1^) (25, 26). However, the final value, which is the only value in the NIST table marked as the “original publication of a measured value,” reports the CO Henry’s Law constant as 7.4 mM·bar^-1^ (25), approximately one order of magnitude higher than the value for H_2_ (this higher value is the CO Henry’s Law constant we used for our calculations). The NIST WebBook does not directly cite the sources for these values, but the site says that the data was compiled by “Rolf Sander.” This presumably refers to Dr. Rolf Sander’s “Compilation of Henry’s law constants (version 5.0.0) for water as a solvent.”

In the Sander reference seven sources are listed as providing a direct measurement of the Henry’s Law constant for CO. Six of these sources report values in the range of 0.71-0.98 mM·bar^-1^, but one source lists a value of 8.0 mM·bar^-1^, similar to the NIST value we used for our calculations (27). This study focuses on the impact of CO at very low aqueous concentrations in the natural environment. The authors show that CO does not properly obey Henry’s Law at high dissolved concentrations. They report that, when the dissolved concentration of CO is sufficiently low, the Henry’s Law constant of CO is an order of magnitude larger than the value which has been widely reported from standard lab measurements using high partial pressures of CO in the headspace (and therefore high dissolved concentrations) (28).

We believe it is highly likely that previous acetogen and gas fermentation researchers have used the lower value of Henry’s Law constant for CO due to the larger number of studies reporting it. However, since acetogens drive the dissolved concentration of gases down when they grow, the Henry’s Law constant for CO measured by Meadows (1974), which was measured using low dissolved concentrations of CO, is likely the most applicable value for gas fermentation. Assuming this higher number is the correct value for the Henry’s Law constant of CO, our experimental data and calculations clearly show that the low growth rate of *C. ljungdahlii* (and all similar acetogens) on CO_2_ and H_2_, relative to CO, is explained by the near order of magnitude decrease in solubility of the electron source (H_2_ vs. CO) during acetogenic growth on CO_2_ compared to CO.

## 3. RESULTS & DISCUSSION

3.1. Fructose drip-feeding improves CO_2_ utilization in fed-batch *C. ljungdahlii* cultures

A potential way to enhance H_2_/CO_2_ utilization is acetogenic mixotrophy (more precisely anaerobic, non-photosynthetic (ANP) mixotrophy, (29)), whereby the cells are fed a sugar or another higher energy-content co-substrate in addition to H_2_/CO_2_. The logic is simple: the higher energy-content substrate provides more energy than the low energy content of gases to achieve better growth (30). The literature suggests that under some conditions and for some acetogens, the presence of another higher-energy content substrate (e.g., a sugar) might inhibit CO_2_ and/or H_2_ utilization (29), but generally, for certain other acetogens, gas utilization under mixotrophic conditions appears not only possible, but also beneficial in achieving higher metabolite yields (9). What is not clear from the literature is the impact of mixotrophy on growth rate, the growth extent and the rate of substrate utilization, all of which appear to depend on the acetogen examined(9). For example, while *C. ljungdahlii* and *C. autoethanogenum* are genetically virtually identical, their mixotrophic metabolism is significantly different (9). The culture medium and mode (batch, fed-batch, continuous chemostat and continuous perfusion) apparently also affect mixotrophic metabolism (9). While Jones et al (9) report good mixotrophic growth of *M. thermoacetica* on gases and fructose, Park et al. (15) suggest that glucose inhibits CO_2_ utilization through some form of “catabolite repression”, which led them to develop a strategy of co-feeding low glucose concentrations in continuous chemostat cultures in order to increase the CO_2_ fixation rate. Although, in contrast to *C. ljungdahlii*, *M. thermoacetica* is a thermophilic acetogen using a large spectrum of carbohydrates and many other high-energy substrates (31), we examined if slow feeding of fructose to *C. ljungdahlii* (which cannot utilize glucose) might benefit gas utilization in fed-batch cultures, which are of translational importance in contrast to chemostat cultures.

We, thus, developed a fed-batch approach aiming to enhance *C. ljungdahlii* growth on CO_2_ by feeding fructose at different rates. For these experiments, we used syringe pumps to drip-feed concentrated fructose solutions into 160mL serum bottles at precise, controlled rates throughout the fermentation. The serum bottle headspace was flushed with an 80/20 mix of H_2_/CO_2_ before cell inoculation, and sufficient H_2_ and CO_2_ were provided by connecting the headspace of each 160-mL bottle to a 500mL serum bottle flushed and pressurized to 10 psig with 80/20 H_2_/CO_2_ mix (Fig. S1A). These larger bottles were used to supply sufficient CO_2_ and H_2_ without over-pressurizing the serum bottles (the syringe pump manufacturer recommended maintaining pressures below 12 psig to avoid inhibiting or damaging the syringe pump). The conditions tested using this fed-batch approach are summarized in Table 1. Each condition was tested using two technical replicates (two separate cultures prepared from the same original *C. ljungdahlii* inoculum). Each condition was provided with 1 g/L fructose in the initial media with the remainder provided through the syringe pump feed. For all experiments presented in this study, we used our *C. ljungdahlii* ptaHALO strain (*Clostridium ljungdahlii* DSM13528 transformed with an anaerobic fluorescent reporter) (16) for its antibiotic resistance to minimize contamination risk.

At all fructose feed rates, fructose levels in the culture were maintained at or below 2 mM for all but the initial timepoint (Figure 1A). The higher fructose levels at the initial timepoint were due to the initial 1 g/L fructose included in the medium and a small amount of the feed solution added at the beginning of the experiment when priming the syringe pump. Approximately 20 mM of ethanol was also present in the starting medium due to the clarithromycin used to maintain the p100ptaHALO plasmid in *C. ljungdahlii* (our clarithromycin stock solutions are prepared in pure ethanol).

**Figure 1:**
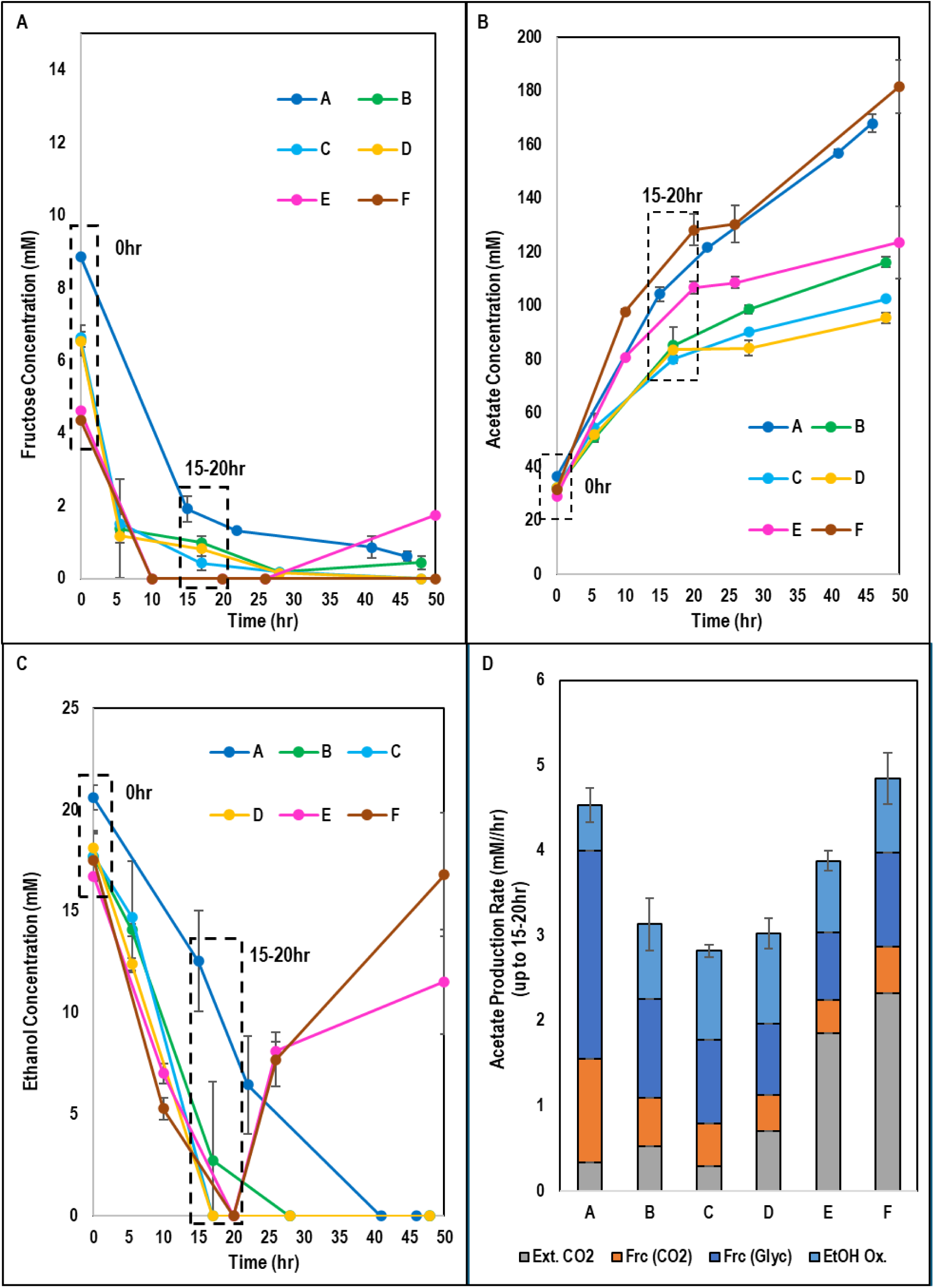
For *C. ljungdahlii* fed-batch fructose feeding experiments. A) Fructose kinetics. B) Acetate kinetics. C) Ethanol kinetics. D) Average acetate production rate (overall and from each carbon source) for the first 15-20hr in each fructose feeding condition. Error bars represent the standard deviation between two technical replicates.

For conditions which had an initial starting cell density of 1 OD_600_ (Table 1; Conditions A-D), the rate and total titer of acetate production over the course of the experiment corresponded to the fructose feed rate. The higher the fructose feed rates the more acetate is produced more quickly (Fig. 1B).

To compare the relative rate of CO_2_ assimilation, we compared the acetate production rate over the first 15-20hr and the source of the produced acetate between these conditions for the first 15-20 hr of fermentation (we only considered up to 15-20 hr because the rate of acetate production slowed considerably for conditions B-E after 15-20 hr) (Fig. 1D. In this case, acetate could come from one of four sources: external CO_2_, CO_2_ released during the decarboxylation of pyruvate to Acetyl-CoA when fructose is processed via glycolysis, acetyl-CoA generated from fructose via glycolysis, and oxidation of the small amount of ethanol present in the initial medium due to the clarithromycin (stock solution dissolved in ethanol) used to maintain the plasmid. The goal of these experiments was to maximize the overall rate and fraction of CO_2_ incorporation, especially external CO_2_.

In condition A, our highest tested fructose feed rate, the cells produced 4.5 mM acetate per hour (over the first 15-20 hrs) and only 7% of this acetate (the sole culture product) came from external CO_2_ (Fig. 1D, Fig. S1C); *C. ljungdahlii* primarily utilized fructose. However, in condition D, which had the same initial starting cell density but the lowest fructose feed rate (nine-fold lower than condition A), 24% of acetate produced came from external CO_2_ (Fig. S1C), and the absolute rate of external CO_2_ fixation was 113% higher than in condition A, although the overall rate of acetate production was lower (Fig. 1D). This suggested that, by optimizing the cell-specific fructose feed rate, the cell-specific CO_2_ fixation rate could also be increased in *C. ljungdahlii* while maintaining low levels of fructose feeding to support biomass production.

Next, in conditions E and F, we tested if, using the same cell-specific fructose feed rate, we could increase the total acetate titer and total rate of external CO_2_ fixation by increasing the initial cell density while maintaining the high fraction of external CO_2_ incorporation. This approach was partially successful, but it gave diminishing returns. The absolute rate of CO_2_ fixation in condition E, which started with twice as many cells as condition D, was 2.6-fold higher than in condition D (Fig. 1D). The overall rate of acetate production was also higher (3.9 mM/hr vs. 3.0 mM/hr), and 48% of this acetate came from external CO_2_, compared to only 24% in condition D (Fig. 1D, Fig. S1C). However, despite starting with twice as many cells as condition E (and therefore four times as many as in condition D), condition F only increased the rate of external CO_2_ fixation 25% compared to condition E (Fig. 1D). The overall rate of acetate production in condition F was also only 25% greater than in condition E, and the percent incorporation of external CO2 into acetate was 48%, identical to condition E (Fig. 1D, Fig. S1C).

These results showed that both the cell-specific and absolute rate of external CO_2_ incorporation could be improved by optimizing the cell density and fructose feed rate. However, these improvements seemed to reach a point of diminishing returns at higher cell densities and, even with these optimizations, no condition produced more than 48% of its acetate from external CO_2_.

We considered two potential explanations for these results. First, we considered that the total amount of cellular energy available from external CO_2_ fixation and the low cell-specific fructose feed rates may have been insufficient to fully satisfy the maintenance energy of the high levels of biomass we tested, especially in conditions E and F, since the cell density, especially in those two conditions, peaked early and then decreased throughout the rest of the experiment (Fig. S1B). Second, we considered that gaseous mass transfer in the serum bottle cultures, not concentration of *C. ljungdahlii* cells, may have become limiting at higher cell densities.

### 3.2. *C. ljungdahlii* CO_2_ fixation is limited by gaseous mass transfer

Minimizing fructose feed rate and accumulation enabled increases in CO_2_ incorporation, but the increases in CO_2_ fixation did not scale proportionally to higher cell densities. This caused us to question if, perhaps, gaseous mass transfer was limiting the CO_2_ fixation rate under the conditions tested, not carbon catabolite repression. To test this hypothesis, we tested the growth and CO_2_ fixation of *C. ljungdahlii* with high fructose concentration (3 g/L) and low fructose concentration (1 g/L) under conditions of high and low mixing. For the low mixing conditions, we grew the cultures in serum bottles on a rotary shaker at 90 RPM (Fig. 2A) (referred to hereafter as the “shaker” condition). For the high mixing conditions, we flipped the 160 mL bottles on their sides and placed them on a tube roller (Fig. 2A, Fig. S2A) (referred to hereafter as the “roller” condition), analogous to the “roller bottle” culture system used for increased oxygen transfer in cell-culture applications (32). Roller bottle reactors favor increased gaseous mass transfer due to formation of a thin liquid film on the bottle walls as it rotates, significantly increasing the available surface area for gas transfer by creating a moving thin film that increases the interfacial gas transport rate and a short distance for the gas to penetrate the liquid due to the very thin film. For all the conditions, each serum bottle was flushed and pressurized to ∼15 psig with an 80/20 mixture of H_2_ and CO_2_, and each bottle was re-flushed and re-pressured to between 15-20 psig after each sampling point to maintain high partial pressure of CO_2_ and H_2_ throughout the experiment. We performed this experiment in biological duplicate twice (four total biological replicates). Results from one set of biological replicates are presented in Figure 2; results from the second set are presented in Figure S3.

**Figure 2:**
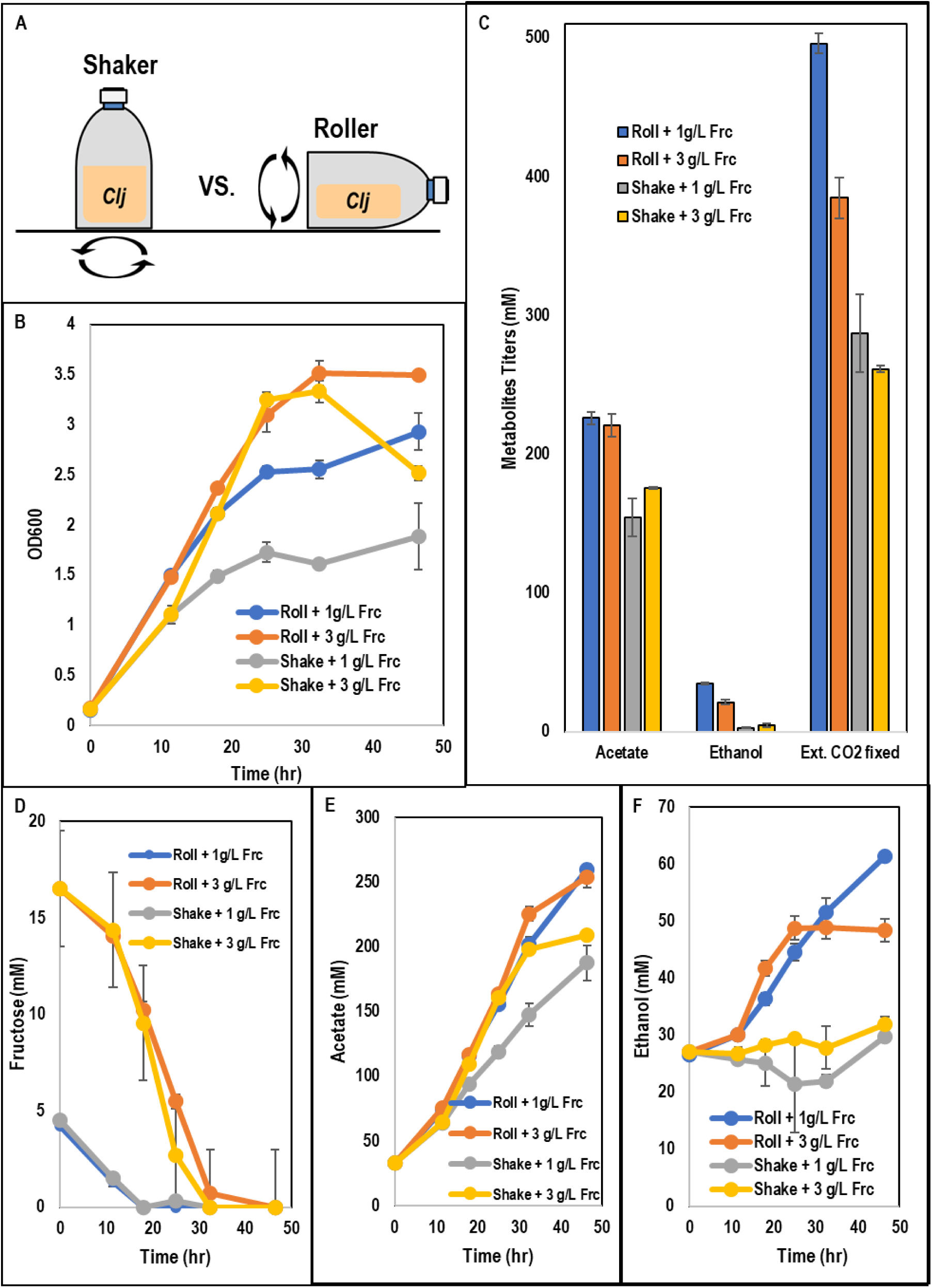
For high/low mixing and high/low fructose batch experiments with *C. ljungdahlii*: A) Schematic of shaker versus roller bottle experimental setup. B) Cell growth kinetics. C) Total metabolite titers. D) Fructose kinetics. E) Acetate kinetics. F) Ethanol kinetics. Error bars represent the standard deviation between two biological replicates. Data from two additional biological replicates are shown in Figure S3.

In this experiment we observed that growth and biomass accumulation were best in the two conditions with high fructose, lower but still strong in the low fructose/roller condition, and weakest in the low fructose/shaker condition (Fig. 2B). Interestingly, we observed a noticeable drop in OD_600_ from 33 hr to 47 hr in the high fructose/shaker condition, corresponding to the time when all fructose was exhausted (Fig. 2B, D). This likely occurred because, once all of the fructose was gone, the cells could not generate energy quickly enough from gases (H_2_/CO_2_) to maintain its biomass, whereas the high fructose/roller condition, which also exhausted fructose around the same time, did not experience a drop in biomass because of the extra energy readily available from gases in the headspace due to increased mixing (thus higher gas transfer rates) in the roller bottles.

In terms of acetate production, both roller conditions substantially outperformed the shaker conditions (Fig. 2C, E), demonstrating the key takeaway from this experiment: gaseous mass transfer is the primary limiting factor for CO_2_ fixation. Interestingly, though acetate represented greater than 85% of total metabolites in all four conditions, ethanol production was much higher in the roller conditions, and it was highest (35 mM, 13% of products) in the low fructose/roller condition (Fig. 2C, F) Significantly, we rarely observe net ethanol production with any culture media made up of H_2_, CO_2_ and fructose or fructose alone. As ethanol production requires more electrons than acetate production, the observed ethanol production suggests that the faster transfer and utilization of CO_2_ and H_2_ provided more electrons to stimulate ethanol production. Total acetate production between the high fructose/roller bottle mode and low fructose/roller conditions was similar (Fig. 2C, E). However, 95% of the total metabolites generated in the low fructose/roller condition came from external CO_2_, whereas 80% of the acetate synthesized in the high fructose/roller condition came from external CO_2_. This translates to 29% more CO_2_ fixed by the low fructose condition in the same amount of time, despite lower biomass accumulation in that condition (Fig. 2B), and thus a much higher rate of cell-specific CO_2_ fixation This result confirms the results of the fed-batch experiment (Fig. 1), demonstrating that the presence of fructose inhibits external CO_2_ utilization. If the presence of fructose did not inhibit external CO_2_ utilization at all, we would expect to observe higher total acetate production in the high fructose/roller condition because that condition would fix the same amount of external CO_2_ as the low fructose condition and then generate additional acetate from the extra fructose present in the growth medium. Collectively, this experiment demonstrates that growth on and fixation of external CO_2_ by *C. ljungdahlii* is limited by both gaseous mass transfer (because both roller bottle conditions make more acetate and ethanol than either shaker bottle condition) and the level of fructose in the medium (because the low fructose/roller condition fixes more external CO_2_ than the high fructose/roller condition).

### 3.3. Enhanced gaseous mass transfer enables *C. ljungdahlii* growth rates and biomass accumulation on CO_2_ and H_2_ comparable to those on fructose, CO, and syngas

We have shown that use of a roller bottle enabled a 66% increase in CO_2_ fixation and 55% increase in biomass accumulation by *C. ljungdahlii* by comparing total external CO_2_ fixed between the low fructose/shake bottle mode and low fructose/roller bottle mode (Fig. 2B, C). Based on that finding we hypothesized that, with sufficient gaseous mass transfer, *C. ljungdahlii* can grow faster and to high cell densities using only CO_2_ and H_2_, similar to its performance on other substrates such as fructose and CO. To test this hypothesis, we performed autotrophic cultures using a similar high (roller bottle mode) and low (shake bottle mode) mixing strategy as in the experiments of Fig. 2, but without fructose.

Four conditions were tested using 20 mL of culture volume in 160-mL serum bottles: batch gas addition in shake bottle mode, batch gas addition in roller bottle mode, fed-batch gas addition in shake bottle mode, and fed-batch gas addition in roller bottle mode. Batch gas addition cultures were flushed with 80/20 H_2_/CO_2_ gas mixture and charged to an initial pressure of ∼20 psig at the beginning of fermentation. Fed-batch gas addition cultures were also flushed with 80/20 H_2_/CO_2_ gas mixture and charged to an initial pressure of ∼20 psig at the beginning of fermentation, but they were also re-flushed and re-pressurized up to ∼20 psig whenever total pressure fell below 15 psig. Fed-batch cultures were meant to improve gaseous mass transport and autotrophic growth relative to batch cultures by maintaining a higher partial pressure throughout the fermentation and by providing more total gaseous substrate throughout the fermentation. We performed two biological replicates each of the batch/shaker/160mL, batch/roller/160mL, fed-batch/shaker/160mL, and fed-batch/roller/160mL conditions.

For additional comparison, an additional condition was tested using 20 mL of culture volume in a 500 mL serum bottle [a higher ratio of gas to liquid volume, (250–20)/20 = 11.5 , vs (160–20)/20 = 7, for the 160-ml bottles] in roller bottle mode. This culture was also performed in fed-batch mode; the culture was pressurized to ∼30 psig with 80/20 H_2_/CO_2_ gas mixture at the beginning of fermentation and re-flushed and re-pressurized to ∼30psig whenever the total pressure dropped below 25 psig. We performed three biological replicates of this fed-batch/roller/500mL culture.

The goal of this additional comparison was to further increase gas transfer by using the largest serum bottles which would fit on our tube roller and the highest gas pressure we were confident these serum bottles could withstand without exploding. Using the 500 mL serum bottles with higher pressure significantly increased gaseous mass transfer relative to the 160 mL serum bottles. This is achieved by increased bottle surface area, which increases the mass transfer coefficient (k_L_a) of the roller bottles, and by being able charge more gas at a time (due to larger bottle volume), which maintains higher total pressure of gas (and therefore higher partial pressures of H_2_ and CO_2_) as gas is used by *C. ljungdahlii* throughout the fermentation (Figure 4B).

The results of these experiments showed improved growth and increased production of acetate from CO_2_ (ethanol was not produced in any condition) in conditions with improved gaseous mass transfer. Without fructose, the differences in growth (Fig. 3A, B) and acetate production (Fig. 4A) between the batch/roller/160mL and batch/shaker/160mL conditions were not statistically significant. Interestingly, both of the batch conditions drove the pressure to sub-atmospheric levels by the end of the fermentation, and the roller condition drove the pressure down more quickly (Fig. 4B). However, the difference between the fed-batch/roller/160mL and batch/roller/160mL (as well as between fed-batch/shaker/160mL and batch/shaker/160mL) was statistically significant, suggesting that the batch cultures were limited by gaseous mass transfer due to decreased partial pressure in the headspace (Fig. 4B). The fed-batch/roller/160mL condition grew faster, accumulated 70% more biomass (an OD_600_ of 2.00 vs. 1.18) and 135% more acetate (275 mM vs. 118 mM) relative to the batch/roller/160mL condition. The fed-batch/roller/160mL condition also outperformed the fed-batch/shaker/160mL condition in terms of growth, growing at a faster rate and accumulating 26% more biomass. These purely autotrophic experiments with the 160mL serum bottles demonstrated that maintaining high partial pressure of the gas mixture (via fed-batch addition of gas) is of similar importance to maximizing mixing. This result is consistent with the gas transfer equation (Eq. 1) (where k_H,gas_, p_gas_, k_L_a, and C^L^_gas_ refer to the Henry’s Law Constant, headspace gas partial pressure, volumetric mass transfer coefficient, and dissolved gas concentration, respectively). In the gas transfer equation, the total gas transfer rate varies proportionally to both mixing (represented by the volumetric mass transfer coefficient “k_L_a”) and headspace gas partial pressure.

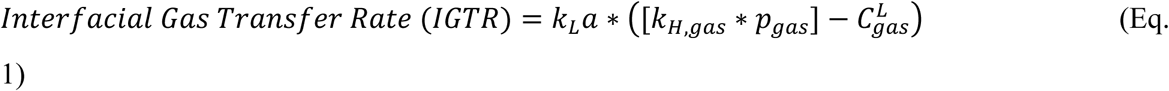

**Figure 3:**
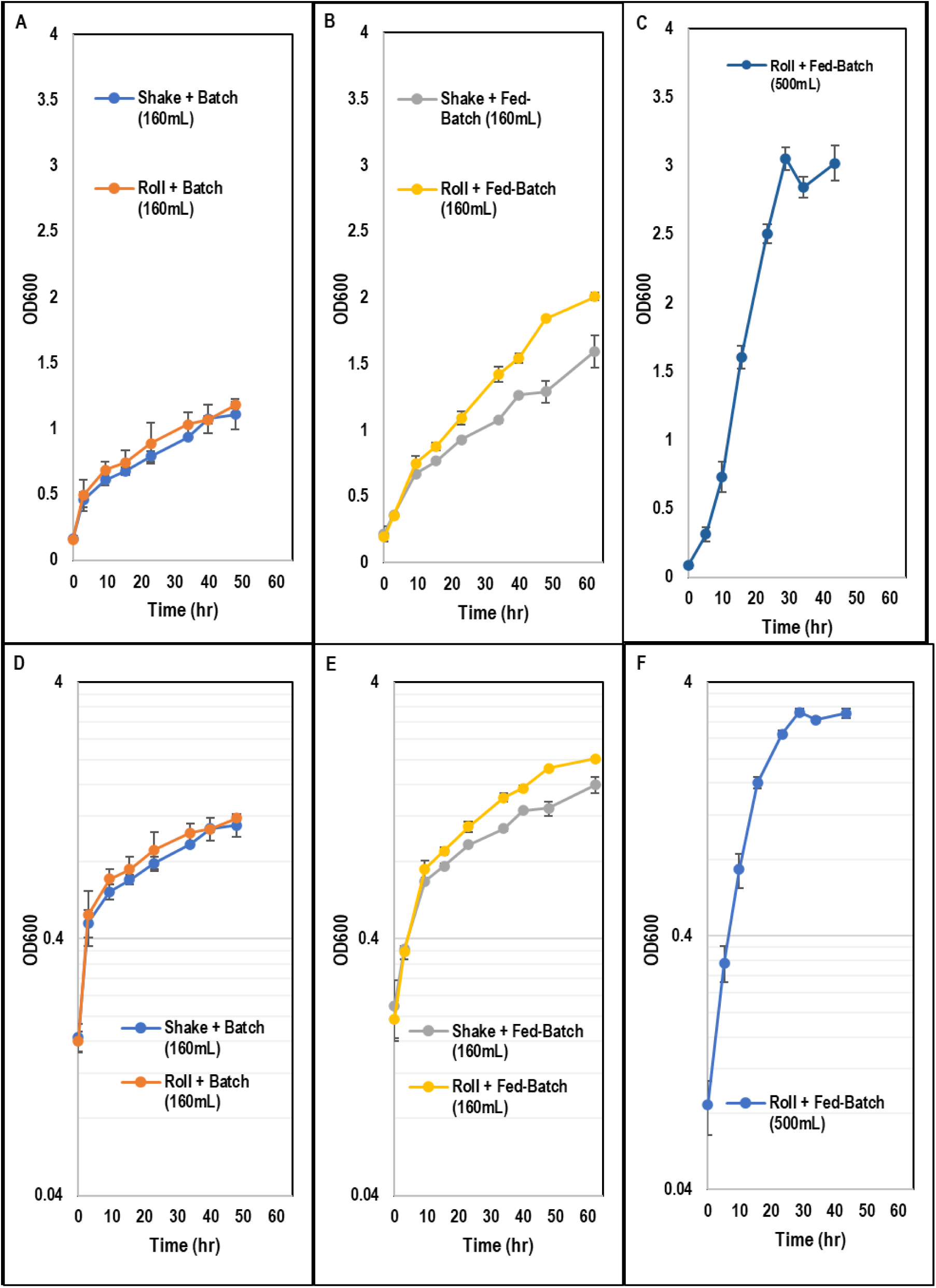
For *C. ljungdahlii* batch and fed-batch growth experiments using only CO_2_/H_2_: A) Cell growth kinetics for batch cultures grown in 160mL serum bottles. B) Cell growth kinetics for fed-batch cultures grown in 160mL serum bottles. C) Cell growth kinetics for fed-batch cultures grown in 500mL serum bottles. D) Log plot of cell growth kinetics for batch cultures grown in 160mL serum bottles. E) Log plot of cell growth kinetics for fed-batch cultures grown in 160mL serum bottles. F) Log plot of cell growth kinetics for fed-batch cultures grown in 500mL serum bottles. For all cultures grown in the 160mL serum bottles, error bars represent the standard deviation between two biological replicates. For cultures grown in the 500mL serum bottles, error bars represent the standard deviation between three biological replicates.

**Figure 4:**
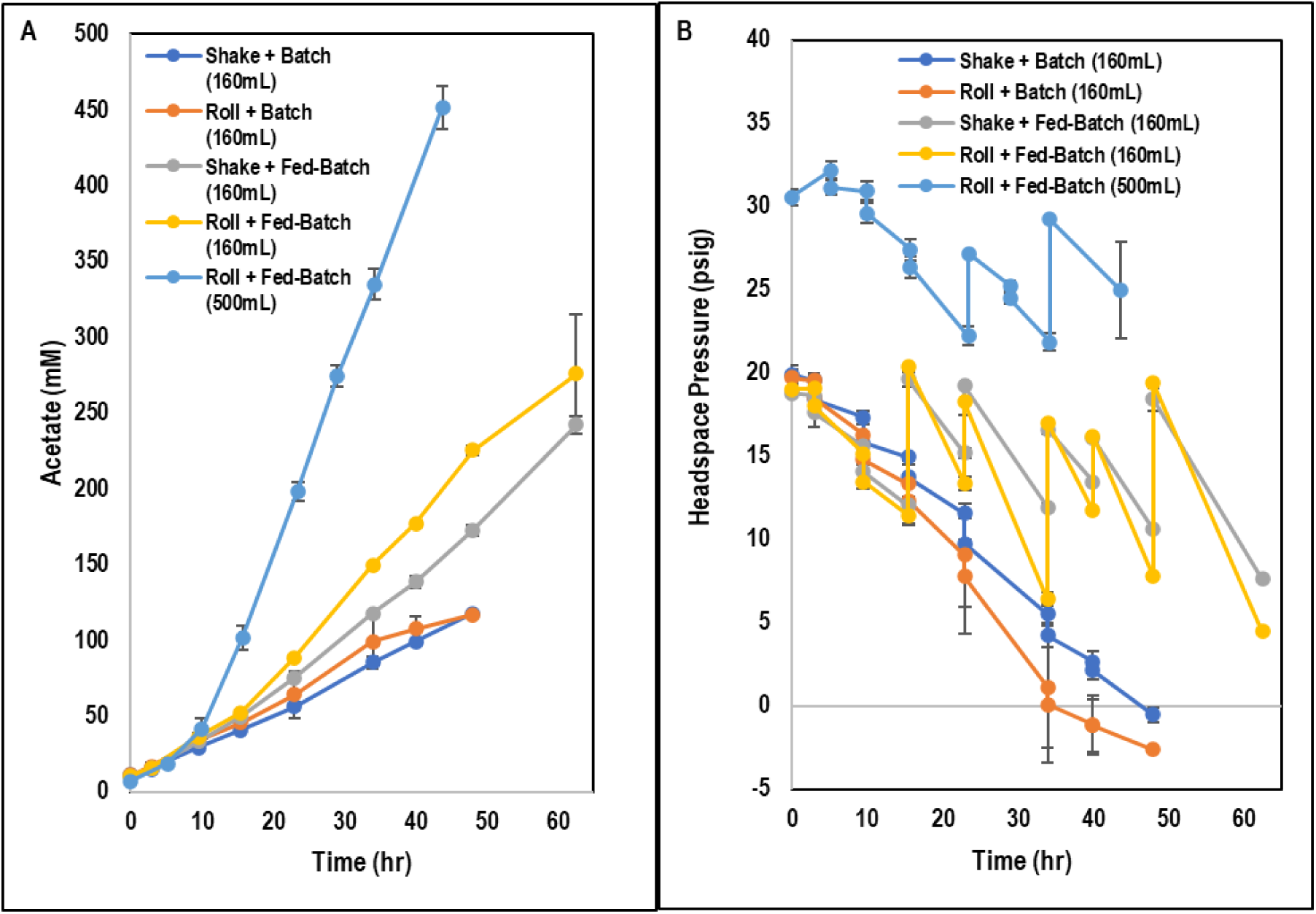
For *C. ljungdahlii* batch and fed-batch growth experiments using only CO_2_/H_2_: A) Acetate kinetics. B) Headspace gas pressure kinetics. C) pH kinetics for batch cultures grown in 160mL serum bottles. D) pH kinetics for fed-batch cultures grown in 160mL serum bottles. F) pH kinetics for fed-batch cultures grown in 500 mL serum bottles. For all cultures grown in the 160mL serum bottles, error bars represent the standard deviation between two biological replicates. For cultures grown in the 500mL serum bottles, error bars represent the standard deviation between three biological replicates.

As hypothesized, the fed-batch/roller/500mL *C. ljungdahlii* fermentations showed major improvements in autotrophic growth and CO_2_ fixation relative to the 160-mL serum bottle fermentations due to the much higher surface area for mixing, higher initial partial pressure (30 psig vs 20 psig), and much larger headspace volume (480 mL vs 140 mL) (as would be expected if acetogenic growth on CO_2_ is limited by gaseous mass transfer). The fed-batch/roller/160mL condition required 63 hrs of growth to reach peak cell density (OD_600_ of 2.01); the fed-batch/roller/500mL culture hit its initial cell density peak of 3.12 OD_600_ after just 29 hr, demonstrating much faster growth (Fig. 3B, C). Over the first 16hr of fermentation, the fed-batch/roller/160mL condition grew at a specific growth rate of 0.097 hr^-1^, a doubling time of 7.1 hr (Fig. 3E). To our knowledge, this would already be the fastest recorded doubling time for *C. ljungdahlii* growing solely on CO_2_ and H_2_ (Table 2). However, over its first 16 hrs of fermentation, the fed-batch/roller/500mL condition grew at a specific growth rate of 0.19 hr^-1^ (doubling time= 3.7 hr) nearly twice as fast as the fed-batch/roller/160mL culture (Fig. 3F).

**Table 2.** Reported *C. ljungdahlii* or *C. autoethanogenum* doubling times when grown on CO, CO_2_/H_2_, and/or fructose.

| Organism | Substrate(s) | Doubling Time (hr) | Source |
| --- | --- | --- | --- |
| *C. ljungdahlii* | CO / H<sub>2</sub> / CO<sub>2</sub><br>(30 / 30 / 30) | 3.6 | Mohammadi et al. (2014) |
| *C. ljungdahlii* | H<sub>2</sub> / CO<sub>2</sub><br>(80 / 20) | 3.7*** | This study (fed-batch/roller/500mL) |
| *C. ljungdahlii* | Fructose + CO / H<sub>2</sub> / CO<sub>2</sub><br>(20 / 10 / 20) | 4.6 | Whitham et al. (2015) |
| *C. ljungdahlii* | Fructose + H<sub>2</sub> / CO<sub>2</sub><br>(80 / 20) | 5.0 | Our lab (unpublished data) |
| *C. ljungdahlii* | Fructose | 6.9 | Oswald et al. (2018) |
| *C. ljungdahlii* | H<sub>2</sub> / CO<sub>2</sub><br>(80 / 20) | 7.1*** | This study (fed-batch/roller/160mL) |
| *C. autoethanogenum* | CO / H<sub>2</sub> / CO<sub>2</sub><br>(50 / 3 / 18) | 7.3 | Valgepea et al. (2017) |
| *C. ljungdahlii* | CO / H<sub>2</sub> / CO<sub>2</sub><br>(23 / 21 / 10) | 9.9 | Oswald et al. (2018) |
| *C. ljungdahlii* | CO / H<sub>2</sub> / CO<sub>2</sub><br>(55 / 30 / 5) | 11.6 | Hermann et al. (2020) |
| *C. ljungdahlii* | H<sub>2</sub> / CO<sub>2</sub><br>(48 / 48) | 28.9*** | Hermann et al. (2020) |
| *C. autoethanogenum* | CO / H<sub>2</sub> / CO<sub>2</sub><br>(2 / 65 / 23) | 16.5 | Heffernan et al. (2020) |
| *C. autoethanogenum* | H<sub>2</sub> / CO<sub>2</sub><br>(67 / 23) | 35.4*** | Heffernan et al. (2020) |
| *C. ljungdahlii* | H<sub>2</sub> / CO<sub>2</sub><br>(80 / 20) | 57.4*** | Klask et al. (2020) |

\*\*\*: uses only CO<sub>2</sub>/H<sub>2</sub>

Interestingly, the semi-log plots of the batch and fed-batch fermentations in 160-mL bottles (Fig. 3D-E) show an initial growth rate comparable to the initial growth rate of the fed-batch fermentation in the 500-mL bottle (Fig. 3F). However, the growth rate in the 160-mL bottles quickly slows as the cell density approaches an OD_600_ of ∼0.8 (Fig. 3E), whereas growth in the 500mL bottles, which has improved mixing and higher gas partial pressure, remains strong up until a cell density of approximately 1.6 (Fig. 3F). This is exactly what we would expect to observe if growth of *C. ljungdahlii* on CO_2_ and H_2_ is mass-transfer limited. The doubling time of 3.7 hr demonstrated here is by far the fastest ever recorded (to our knowledge) for *C. ljungdahlii* or *C. autoethanogenum* using only CO_2_ and H_2_ and it is nearly identical to the fastest doubling time ever recorded (to our knowledge) for *C. ljungdahlii* of 3.6 hrs, a result which was achieved growing *C. ljungdahlii* in well-mixed, horizontal, 160mL serum bottles in PETC medium under a syngas headspace which included 30% CO (Table 2). Notably, our results (as well as the best results achieved on CO) were achieved in serum bottles without pH control (Fig. S4A-C) or constant gas addition (Fig. 4B), not a well-controlled bioreactor. These results strongly suggest that, with sufficient gaseous mass transport, *C. ljungdahlii* grows just as well on CO_2_ and H_2_ as on any other substrate.

This improvement in autotrophic growth rate also translated to robust acetate production. Whereas the fed-batch/roller/160mL culture required 63 hrs to produce 276 mM of acetate, the fed-batch/roller/500mL condition produced 452 mM of acetate in only 44 hrs, corresponding to a 2.3 times increase in acetate productivity (and CO_2_ fixation). Even though biomass accumulation in the 500-mL bottle condition plateaued between 29 and 44 hrs, acetate production continued at essentially a constant rate, suggesting that the cells were still highly active through the end of the fermentation. Despite manual pH adjustments at every timepoint, pH varied wildly throughout all of these experiments (Fig. S4A-C) from above 6.0 to below 5.0 with little apparent impact on growth or the acetate production rate.

### 3.4. Cellular energy balance and gas transport calculations suggest that low *C. ljungdahlii* growth rate and biomass accumulation on CO_2_/H_2_ relative to CO is due to low H_2_ solubility in culture media

If (assuming sufficient gaseous mass transfer) *C. ljungdahlii* can grow just as quickly on CO_2_ and H_2_ as on CO or fructose, why do so many studies indicate the opposite (Table 2)? Also, it is not clear why *C. ljungdahlii* produces exclusively acetate when grown on CO_2_/H_2_ instead of the ethanol typically produced when grown on CO. Leaving aside fructose and considering only CO_2_/H_2_ and CO (gas fermentations), we considered the impact of interfacial transport [IGTR, Equation (1)]. Assuming the same mixing intensity (same k_L_a) and partial pressure (p_gas_), IGTR is affected by the Henry’s Law constant (k_H,gas)_. The k_H_ for CO_2_ is 34 mM*bar^-1^ (33), and 7.4 mM*bar^-1^ for CO (25), meaning that the solubility of CO_2_ is 4.6-fold higher than CO. However, crucially, k_H_ for H_2_ is only 0.78 mM*bar^-1^, 9.5-fold lower than CO (26). This near order-of-magnitude difference in the solubilities between H_2_ and CO adequately explains the difference in growth and product production of *C. ljungdahlii* (& *C. autoethanogenum*) on CO_2_ and H_2_ compared to CO. This can be demonstrated by combining the well-established metabolic pathway energetics for growth on CO and CO_2_/H_2_, gas transfer, and gas solubility in the cellular energy balance.

If we assume that CO_2_ and H_2_ are the only substrates for growth, we can equate the microbial growth equation with the gaseous mass transfer rate of H_2_ according to the IGTR (Eq. 1) multiplied by the yield coefficient of ATP per H_2_ fixed to derive Equation 2.

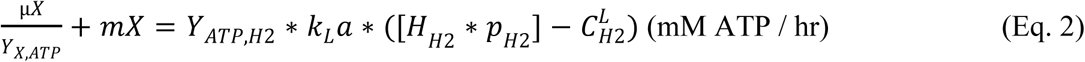

μ represents the microbial specific growth rate (units of hr^-1^), “X” represents the *C. ljungdahlii* biomass concentration using any appropriate unit (we will use gDCW*L^-1^), “Y” represents the molar yield coefficient of millimolar amounts ATP generated per millimole of H_2_ used (mM ATP/mM H_2_) , and “m” represents the cellular maintenance energy coefficient, using units consistent with µ, X and Y [in this case gDCW*L^-1^*hr^-1^*(mM ATP)^-1^). We consider here only H_2_ transport because CO_2_ is 44-fold more soluble than H_2_, and 2 mole H_2_ are required per mole CO_2_ (for acetate production via the WLP), meaning that H_2_ transport will be the limiting factor for acetogenic growth on CO_2_ and H_2_ so long as the CO_2_ partial pressure is greater than 1/88^th^ (2*44 = 88) of the H_2_ partial pressure.

For Equation 2, we can set the dissolved H_2_ concentration to zero, since this term will correspond to the H_2_ threshold (“H_2_ threshold” refers to the minimum dissolved H_2_ concentration at which the organism can utilize H_2_ (34)) for *C. ljungdahlii* (2.22 x 10^-3^ mM H_2_) (35), which is negligible compared to the maximum possible driving force (k_H,H2_*p_H2_) at typical H_2_ partial pressures used in gas fermentation. For a typical fermentation in our lab using 20 psig (2.4 bar) of an 80% H_2_ gas mix: k_H,H2_*p_H2_ = k_H,H2_*y_H2_*P = (0.78 mM*bar^-1^)*0.8*(2.4 bar) = 1.5 mM H_2_. We thus write Equation 3.

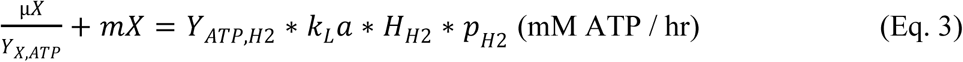

The left-hand side of Equation 3 now represents the rate of ATP generation *C. ljungdahlii* requires for growth and maintenance at a given cell density and specific growth rate, and the right-hand side represents the maximum rate at which cellular energy can be generated from the H_2_ and CO_2_ in the headspace.

From this expression, we see that the total amount of cellular energy available to *C. ljungdahlii* is directly proportional to the stoichiometric ATP yield of H_2_ (from the WLP), the mixing intensity (represented by the volumetric mass transfer coefficient, k_L_a), the Henry’s Law constant for H_2_, and the partial pressure of H_2_ (proportional to the total pressure and/or molar fraction of H_2_). Though we have used H_2_ as an example, Equation 2 is equally applicable to growth with CO, CO_2_, or any other insoluble gaseous substrate when that substrate is the sole source of cellular energy. Equation 3 is similarly applicable so long as the substrate gas threshold is negligible compared to the maximum driving force (k_H,gas_*p_gas_).

Now, we can substitute the different Henry’s Law constants and ATP yields for CO and H_2_ into Equation 3 and compare the output (Table 3). This result shows that, if we assume the same mixing intensity (same k_L_a), the same partial pressure of the gaseous substrate, and assume production of acetate as the sole product; 17.5 times more cellular energy is available to *C. ljungdahlii* from CO compared to H_2_ (and CO_2_) due to the near order of magnitude higher solubility of CO and 1.75-fold higher ATP yield.

**Table 3.** Determining maximum ATP generation rate and cell density for *C. ljungdahlii* as functions of the volumetric mass transfer coefficient (k_L_a) and gas substrate partial pressure.

| Gas Substrate(s) | Target Product | Stoichiometric Equation<br>(derived using WLP stoichiometry from Schuchmann & Muller 2014) | Soluble Product Carbon Recovery | $Y_{ATP/substrate}$<br>(mol ATP per mol CO or H<sub>2</sub> for target product) | Henry's Law Constant<br>(mM·bar<sup>-1</sup>) | Maximum ATP Generation Rate<br>(mM ATP·hr<sup>-1</sup>)<br>(from Eq. 3) | Maximum Cell Density<br>(gDCW·L<sup>-1</sup>)<br>(from Eq. 4) | Maximum ATP Generation Rate<sup>a</sup><br>(normalized) | Maximum Cell Density<sup>b</sup><br>(normalized) |
| --- | --- | --- | --- | --- | --- | --- | --- | --- | --- |
| CO | Acetate | 4 CO →<br>1 Acetate + 2 CO<sub>2</sub> + 1.13 ATP | 0.50 | 0.28 | 7.4 | 2.1 ($k_La \cdot P_{CO}$) | 2.1 ($k_La \cdot P_{CO} \cdot m$) | 17.50 | 17.50 |
| CO | Ethanol | 6 CO →<br>1 Ethanol + 4 CO<sub>2</sub> + 0.63 ATP | 0.33 | 0.11 | 7.4 | 0.81 ($k_La \cdot P_{CO}$) | 0.81 ($k_La \cdot P_{CO} \cdot m$) | 6.75 | 6.75 |
| H<sub>2</sub>, CO<sub>2</sub> | Acetate | 2 CO<sub>2</sub> + 4 H<sub>2</sub> →<br>1 Acetate + 0.63 ATP | 1.00 | 0.16 | 0.78 | 0.12 ($k_La \cdot P_{H_2}$) | 0.12 ($k_La \cdot P_{H_2} \cdot m$) | 1.00 | 1.00 |
| H<sub>2</sub>, CO<sub>2</sub> | Ethanol | 2 CO<sub>2</sub> + 6 H<sub>2</sub> →<br>1 Ethanol + 0.20 ATP | 1.00 | 0.033 | 0.78 | 0.026 ($k_La \cdot P_{H_2}$) | 0.026 ($k_La \cdot P_{H_2} \cdot m$) | 0.22 | 0.22 |

<sup>a</sup>: These values were obtained by dividing each entry in the “Maximum ATP Generation Rate” column by the entry for production of acetate from H<sub>2</sub>, CO<sub>2</sub> to show its relative value to that entry when controlling for mixing ( $k_La$ ) and gas partial pressure.
<sup>b</sup>: These values were obtained by dividing each entry in the “Maximum ATP Generation Rate” column by the entry for production of acetate from H<sub>2</sub>, CO<sub>2</sub> to show its relative value to that entry when controlling for mixing ( $k_La$ ) and gas partial pressure.

Using this approach, we can also compare the energetic requirements to produce different products, such as ethanol, from CO or H_2_/CO_2_. Indeed, this type of comparison clearly shows why making only ethanol from CO_2_ and H_2_ using *C. ljungdahlii* is so difficult. Not only is energy yield per ethanol generated lower than the energy yield per acetate generated (0.2 ATP vs 0.63 ATP; Table 3), producing 1 mol of ethanol requires 6 mol of H_2_, compared to only 4 mol of H_2_ per mol acetate, meaning that the overall yield coefficient of ATP per H_2_ when making ethanol is nearly 5-fold lower than when making acetate (0.033 vs. 0.16; Table 3). In this sense, the inability of *C. ljungdahlii* (and similar acetogens) to make ethanol instead of acetate from CO_2_ and H_2_ is driven by bioenergetics. However, this energy yield disparity can be readily overcome when *C. ljungdahlii* (and similar acetogens) grow on CO due to its nearly 10-fold higher solubility and the more than 3-fold higher energy yield for producing ethanol from CO. If we compare the cellular energy available to *C. ljungdahlii* making only acetate from H_2_/CO_2_ compared to *C. ljungdahlii* making only ethanol from CO, the maximum rate of ATP generated for the ethanol-from-CO case is still 6.75 times higher than the acetate-from-H_2_/CO_2_ case (Table 3), even though the energy yield of acetate from H_2_/CO_2_ is higher than the energy yield of ethanol from CO, because of the difference in solubility between CO and H_2_.

These calculations explain the vast difference in cellular performance between *C. ljungdahlii* and *C. autoethanogenum* when grown on CO_2_ and H_2_ compared to CO, as well as the immense difficulty in adapting or engineering acetogens to produce anything other than acetate from CO_2_ and H_2_. Different bioenergetics between use of CO and CO_2_ and production of ethanol and acetate play a role, but the primary issue is physical chemistry, that is, different gas solubilities.

### 3.5. Maximum cell density in gas fermentation is directly proportional to the solubility of the electron source (H_2_ or CO) due to cellular maintenance energy requirements

For large scale microbial bioprocesses, cell retention is generally used to build cell density and achieve the high volumetric productivities required for economic feasibility. Under conditions of high cell retention in a continuous reactor, microbial cells primarily operate in maintenance metabolism, not growth metabolism. Therefore, we can set the cellular growth term from Eq. 3 equal to zero to assume 100% cell retention (though, in practice, a 5-10% cell bleed is used to remove dead cells and induce some cell growth) (Eq. 4). Then, we see that the maximum steady-state cell density by an acetogen achievable under conditions of full cell retention (i.e. a retentostat) is directly proportional to the total available cellular energy from H_2_ and equal to the total available cellular energy divided by the maintenance coefficient (Eq. 4).

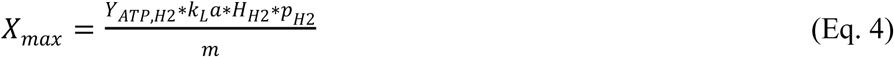

Therefore, we can restate the earlier comparison for *C. ljungdahlii* growing on either CO or CO_2_/H_2_ as follows: in a steady state retentostat *C. ljungdahlii* will be able to accumulate and maintain 17.5 times more biomass when growing on CO compared to CO_2_ and H_2_ (assuming equal partial pressures of CO and H_2_, equivalent mixing, and production of acetate as the sole product) (Table 3). Even if we assume complete ethanol production on CO and complete acetate production on CO_2_ and H_2_, maximum cell density on CO will still be 6.75-fold greater than on CO_2_ and H_2_ (Table 3) (equivalent to the difference in maximum rate of ATP generation calculated earlier).

This result explains why it is much more difficult to grow acetogens such as *C. ljungdahlii* and *C. autoethanogenum* to high cell densities on only H_2_/CO_2_ compared to gas mixtures which contain CO (12, 13); the maximum cell density depends on the maximum rate of energy generation divided by the maintenance coefficient. Although we used the example of cell retention in a continuous reactor, similar logic applies in a batch culture. For a given gas substrate, gas substrate partial pressure, and mixing environment, *C. ljungdahlii* (and similar acetogens) can generate ATP at a certain maximum rate. When they start out at low cell density, most of this energy can be directed towards growth. However, as the cell density rises, the energy required to maintain those cells (which is equal to m*X) also increases until it is equal to the maximum rate of ATP generation, at which point the cells stop growth (or at least net growth). Thus, for a given maintenance coefficient, the maximum cell density will be proportional to maximum rate of ATP generation. For acetogens (as we have discussed at length) this maximum rate of ATP generation, and thus the maximum cell density, will be much higher when using the WLP to grow on CO compared to using the WLP to grow on H_2_/CO_2_ (Table 3).

## 4. CONCLUSION

In this study, we present data suggesting that, though other carbon sources (such as fructose) can suppress CO_2_ utilization, the low solubility of H_2_ is the primary limiting factor for CO_2_ fixation by *C. ljungdahlii* and similar acetogens. How can this issue be overcome to achieve economically viable, CO_2_-negative gas fermentation? Obviously, developing industrial-scale reactors with major improvements in gaseous mass transfer would help. However, mixing to maximize interfacial gas transfer in bioreactors is a well-developed art, and, even if a transformative breakthrough were to take place to further increase k_L_a, CO fermentation would continue to outperform CO_2_ and H_2_ fermentation by approximately an order of magnitude. As clearly demonstrated in the equations of Table 3, when CO is used as carbon and electron substrate, at least 50% (for acetate production, 67% for ethanol production) of it is oxidized and released as CO_2_, thus defeating the goal of carbon-negative production of chemicals.

Fortunately, other avenues exist. We have recently shown that mixotrophic coculture of the acetogen *C. ljungdahlii* with *Clostridium acetobutylicum*, a solventogenic carbohydrate fermenting organism, enables CO_2_-negative mixotrophic fermentation and production of isopropanol as the major product (19). When grown on glucose, *C. acetobutylicum* produces large quantities of H_2_. In coculture *C. ljungdahlii* can use this aqueous H_2_ directly from the liquid medium as it is generated by *C. acetobutylicum* (and may also obtain electrons from *C. acetobutylicum* via direct cell-to-cell contact). Thus, a large fraction of the electrons and H_2_ used to fix CO_2_ by *C. ljungdahlii* in the coculture does not have to overcome the gas-to-liquid mass transfer barrier. Another major benefit of this approach is that *C. acetobutylicum* can upgrade the acetate produced by *C. ljungdahlii* into a diverse range of products, including acetone, isopropanol, and C4 acids and alcohols (17). An additional option would be to include chain elongating organisms, such as *Clostridium kluyveri*, to utilize acetate produced from an acetogen and alcohols produced by a solventogen for CO_2_-negative, mixotrophic production of C4-C8 chemicals (36, 37).

An additional promising approach is to use soluble electron carriers, such as formate or methanol, in place of or in addition to H_2_ to reduce CO_2_. Several acetogens, including *A. woodii*, can assimilate methanol through the Wood-Ljungdahl Pathway (38, 39), and both *A. woodii* and *C. ljungdahlii* (and many other acetogens) can assimilate formate (40, 41). Methanol can be synthesized from methane or sourced renewably (42), and formate has been proposed as a promising storage and transport mechanism for renewable H_2_ via electrochemical reduction of CO_2_ (43). If acetogens can be adapted or engineered to utilize highly soluble methanol or formate alongside or in place of H_2_ and/or CO_2_, the mixing requirements for gas fermentation at scale could be significantly decreased (or even eliminated), simplifying and transforming the design and scaling of economically viable, CO_2_-negative fermentation. Such an approach could also be combined with the aforementioned coculture approach to enable scalable, mixotrophic, CO_2_-negative fermentation to a wide range of products.

## SUPPLEMENTAL MATERIAL

**Supplementary Figures (Word Document)** (Figures S1-S4)

## ACKNOWLEDGEMENTS

This work was supported by an ARPA-E project under contract AR0001505. N.B.W. was supported in part by a U.S. Department of Education GAANN Fellowship under grant P200A210065.

E.T.P. and N.B.W. conceived the project. E.T.P. and N.B.W. designed the experiments. N.B.W. and P.A.B. performed all experiments. E.T.P. and N.B.W. analyzed the data and wrote the manuscript.

---

**Figure S1:**
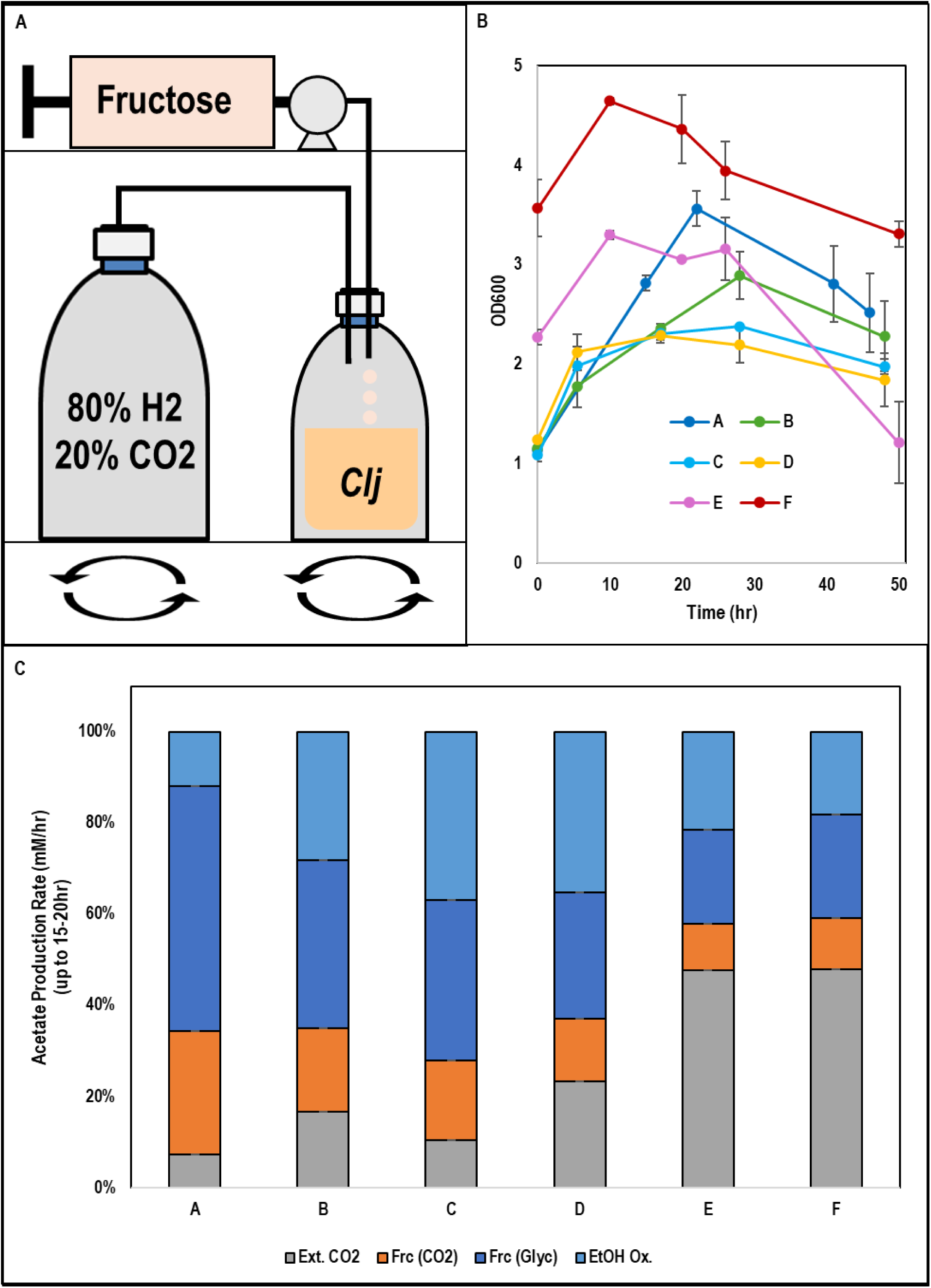
For *C. ljungdahlii* fed-batch fructose experiments: A) Schematic of experimental apparatus. B) Cell growth kinetics. C) Percent of total acetate production rate from each carbon source over the first 15-20hr of fermentation. Error bars represent the standard deviation between two technical replicates.

**Figure S2:**
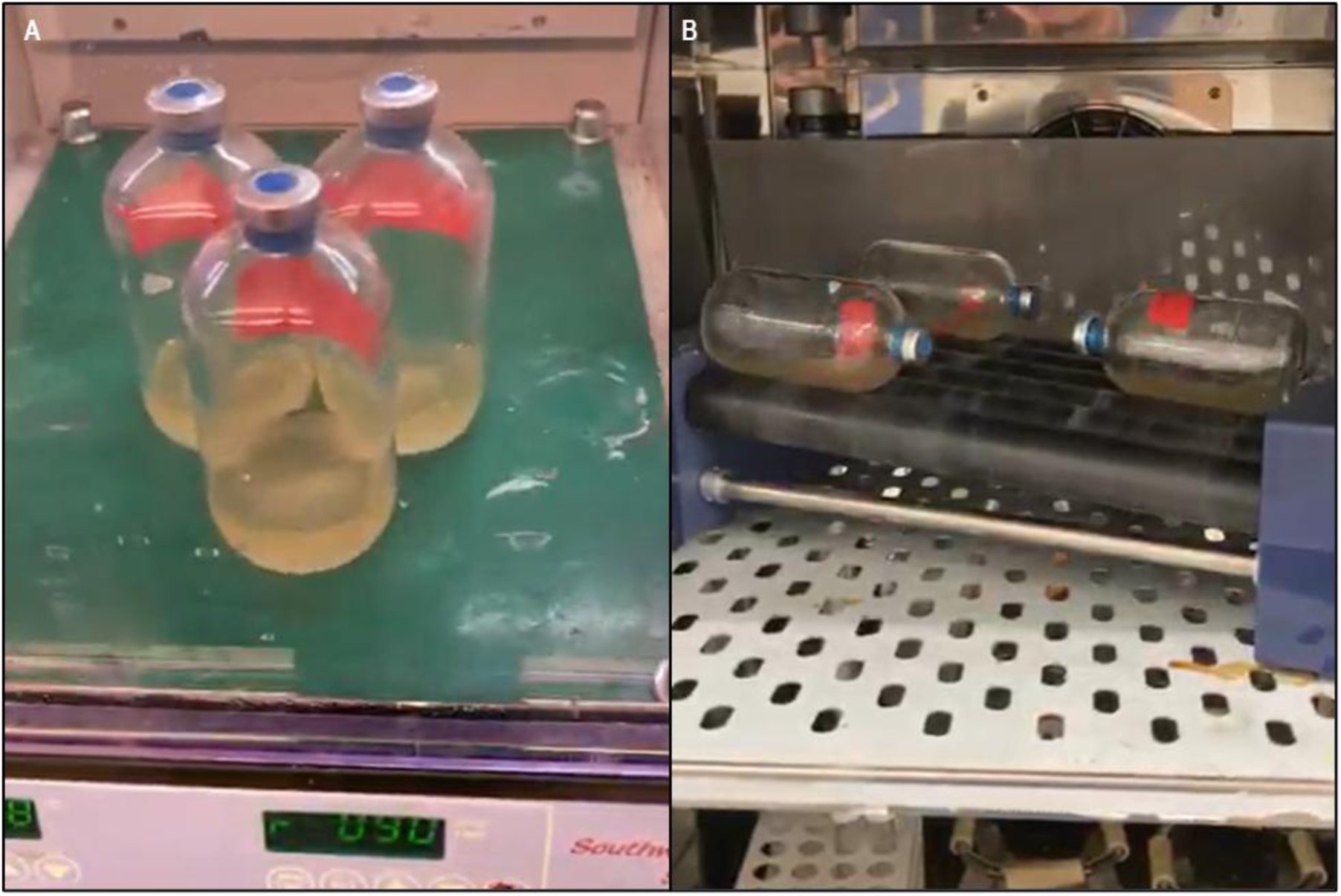
A) Picture of *C. ljungdahlii* cultures grown on rotary shaker. B) Picture of *C. ljungdahlii* cultures grown in roller bottles on tube roller.

**Figure S3:**
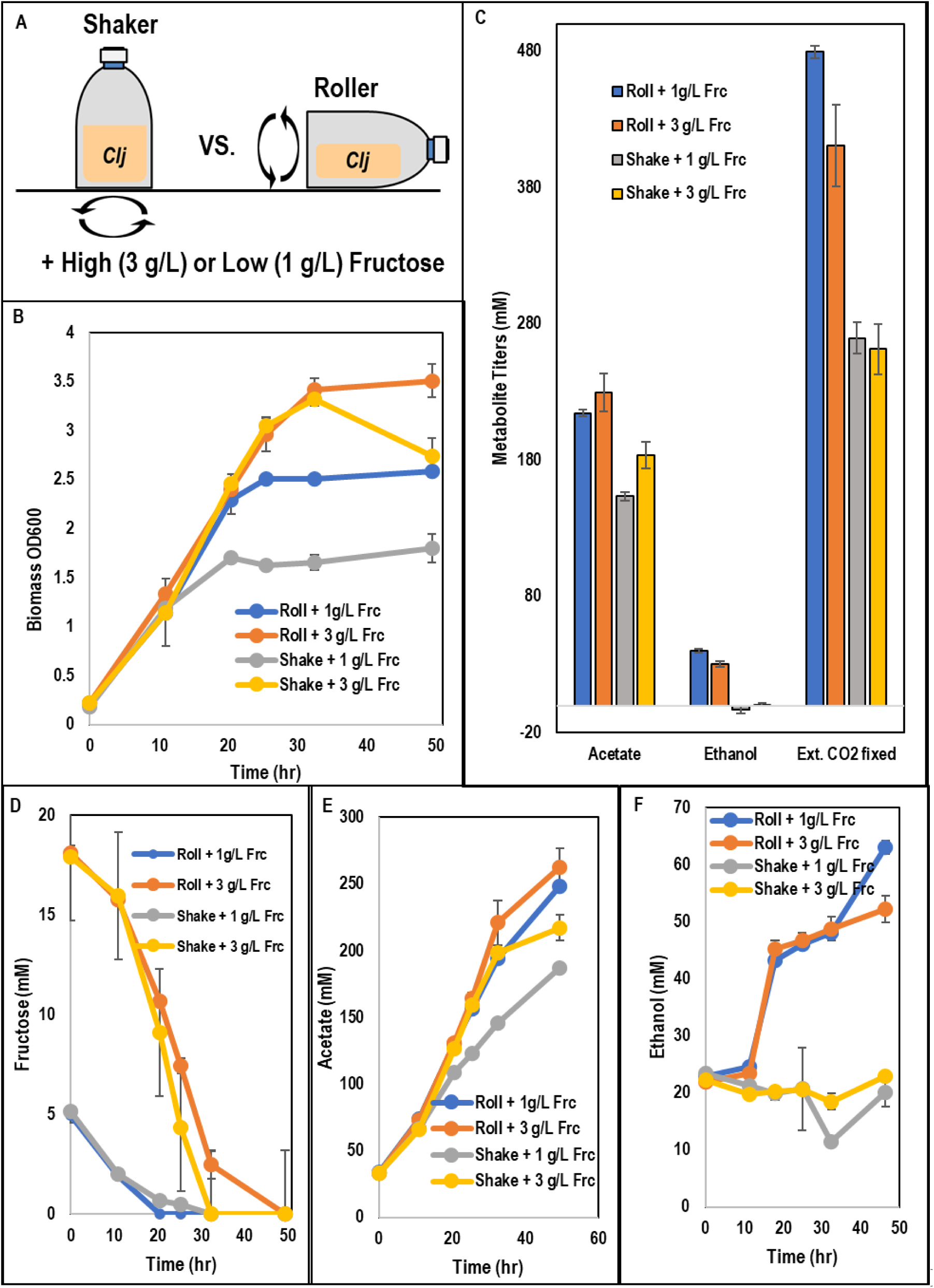
For high/low mixing and high/low fructose batch experiments with *C. ljungdahlii*: A) Schematic of shaker versus roller bottle experimental setup. B) Cell growth kinetics. C) Total metabolite titers. D) Fructose kinetics. E) Acetate kinetics. F) Ethanol kinetics. Error bars represent the standard deviation between two biological replicates. Data from two additional biological replicates are shown in Figure 2.

**Figure S4:**
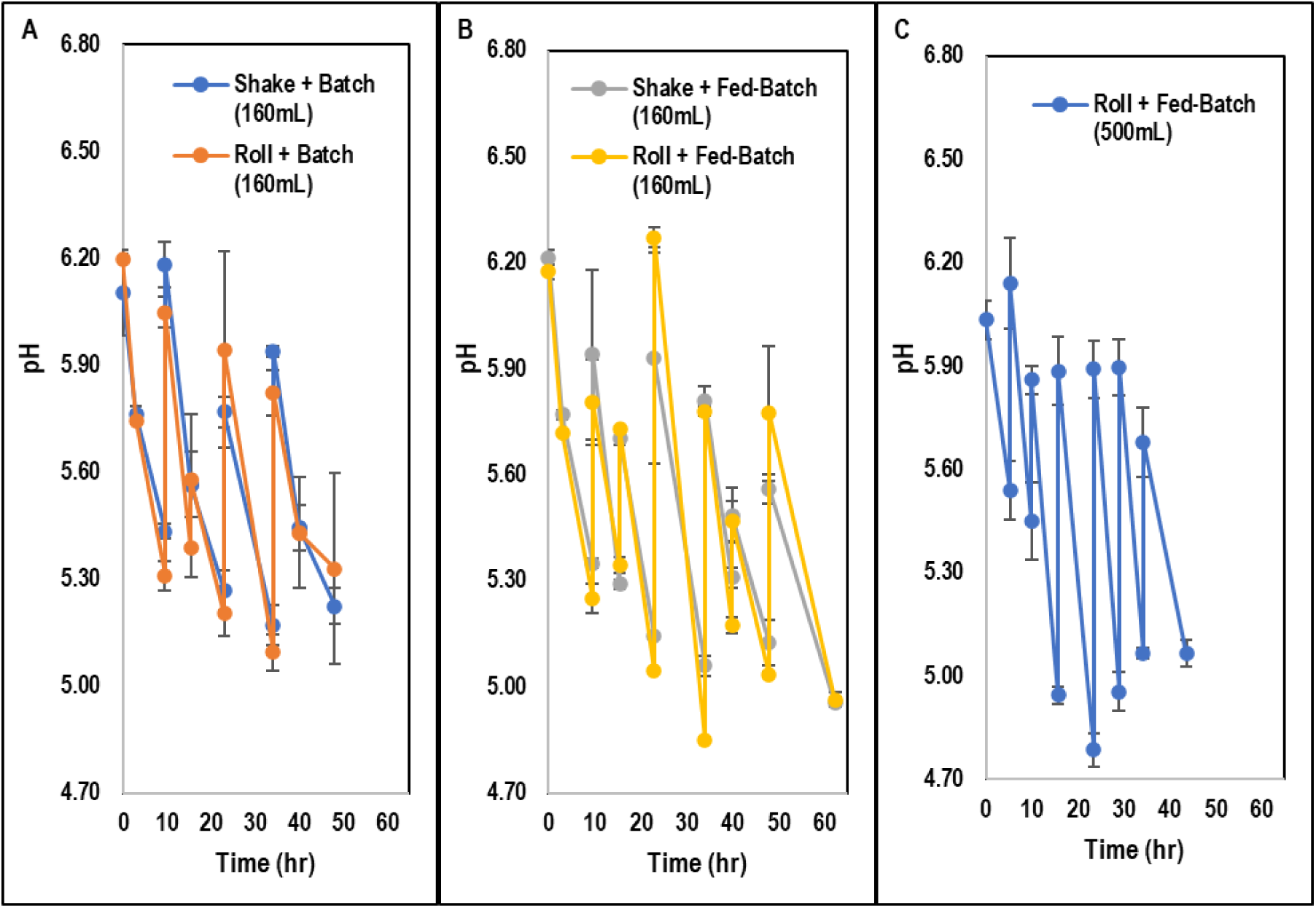
For C. ljungdahlii batch and fed-batch growth experiments using only CO2/H2: A) pH kinetics for batch cultures grown in 160mL serum bottles. B) pH kinetics for fed-batch cultures grown in 160mL serum bottles. C) pH kinetics for fed-batch cultures grown in 500 mL serum bottles. For all cultures grown in the 160mL serum bottles, error bars represent the standard deviation between two biological replicates. For cultures grown in the 500mL serum bottles, error bars represent the standard deviation between three biological replicates.

